# Searching for Jouvence SnoRNA in the Horse Genome

**DOI:** 10.1101/2024.11.21.624650

**Authors:** Mandy Peffers, Guus van den Akker, Emily Clarke, Yongxiang Fang

## Abstract

In humans, various pathologies have been associated with snoRNAs. Others have shown that the expression of a non-canonical snoRNA Jouvence is involved in lifespan determination in relation to gut homeostasis. As snoRNAs are evolutionary conserved, both structurally and functionally, a jouvence orthologue has been identified in humans. This study aimed to locate the Jouvence snoRNA in the horse genome. Using our previous snoRNA data in ageing equine cartilage along with equine genome data we identified a putative equine Jouvence snoRNA gene. ECABCGRLG0000000730 was the primary candidate horse Jouvence snoRNA. The expression of equine Jouvence snoRNA was increased in ageing equine cartilage.

## Introduction

The project aims to locate the Jouvence snoRNA in the horse genome, which was reported in drosophila [1], mouse and human [2]. The study reported the expression of this snoRNA impacts the lifespan of drosophila. Whilst overexpression of Jouvence, which appeared to be a non-canonical H/ACA snoRNA, represented a new tool to fight against the deleterious effects of ageing, while inversely, its knockdown could represent a new approach in cancer treatment. This current study aimed to identify whether Jouvence snoRNA was also located in the horse genome by using our small RNA sequencing data generated from a previous project on ageing equine chondrocytes[3] and the equine genome.

## Methods

The initial processing and quality assessment of the sequence data, genome reference data preparation and reads alignment were conducted previously. The annotation information collected from different sources plus the outcome from the detection of novel snoRNA are the starting point for locating the Jouvence snoRNA in the horse genome. Read alignment information, which has already been generated and previously reported, can also be useful for this study. The relevant processes and summarised are included as below.

### Alignment to reference sequences

The genome reference sequence and annotations used for alignment were downloaded from NCBI. The link for the DNA reference sequence is: ftp://ftp.ncbi.nlm.nih.gov/genomes/all/GCF/002/863/925/GCF_002863925.1_EquCab3.0/GCF_002863925.1_EquCab3.0_genomic.fna.gz

The link of the annotation gff is: ftp://ftp.ncbi.nlm.nih.gov/genomes/all/GCF/002/863/925/GCF_002863925.1_EquCab3.0/GCF_002863925.1_EquCab3.0_genomic.gff.gz

Annotation information of gene features are also linked to other databases:

ftp://ftp.ebi.ac.uk/pub/databases/Rfam/14.0/

ftp://mirbase.org/pub/mirbase/CURRENT/genomes/eca.gff3

The information in Dbxref filed from genome gff was used to link to the above databases to extract more commonly used gene name. Alignment of reads was carried out using above genome reference. TopHat version 2.1.0 [4] was used as the alignment tool with the option “-g 1”, which instructs the software to report the best hits, or randomly select one if there are more than 1 hits that are equally best.

### Counting aligned reads to features

Reads aligning to the reference genome sequences were counted according to the gene features that they mapped to, as defined in the GTF files, using HTSeq-count version 0.6.1p1 [5].The features whose biotype belonged to the gene categories such as miRNA, snoRNA, ncRNA etc, were extracted to generate a categorised gene reads-count table.

### Approaches to characterise putative Jouvence snoRNA

Low identity (around 50%) was observed in global alignments of the Jouvence human and mouse snoRNA on fly’s Jouvence snoNRA sequence [1]. This suggested that identifying the location of this snoRNA in horse would be difficult by homology searches alone. Further, though very likely, it is not certain at this point, whether this snoRNA exists in the horse genome. Therefore, the important information used to characterise this RNA were its length, type of motif box, secondary structure. This information was be used to identify raw candidate Jouvence snoRNAs, but there would likely be many false-positives and these criteria alone would not be sufficient to accurately and confidently identify the snoRNA. For this reason, expression information was also important to filter out candidates; specifically, all candidate genomic regions identified using the above criteria were assessed to determine whether they had RNA read data aligned to them, with a focus on whether (a) the full (or nearly full) length of the candidate was covered by read data and (b) there was reasonable depth of coverage (to eliminate noise). Those that do not pass these criteria will be filtered out.

Based on the above overview, this study was carried out using two different input sets. One set contained 863 novel snoRNA which were present in the horse genome annotation set [3], but have no further functional annotation (named as N-S set). The second set included 5187 sequences which were extracted from the horse genome on a pattern constructed based on characteristics of the drosophila Jouvence snoRNA. This was compared it to the corresponding one in the human genome (named as P-E set).

### Detection of jouvence snoRNA from the two sets

Primary selection (stage 1)was undertaken by obtaining read alignment information for the sequences in the subject sequence sets (using tophat, salmon). Then the read counts were calculated for sequences in the subject sequence sets and sequences removed which were not aligned to by any reads Then the overlap between extracted sequences and exon features was checked, and excluding if sequences overlapped with any non-snoRNA exons. The remaining sequences which were H/ACA class snoRNAs were checked and, excluding those reported as not detected by snoReport [6] were excluded. Sequences with length < 139 pb or > 155bp were removed.

The selection was then refined by checking whether the majority of a sequence was covered by reads (stage 2). 95%+ of sequence length was proposed as the lower boundary. The secondary structure was checked of remaining H/ACA snoRNAs with snoReport, selecting the one with the highest similarity in secondary structure with Drosophila Jouvence snoRNA.

Next a across species comparison was undertaken using pairwise global alignment of Drosophila Melanogaster, Human and detected Horse Jouvence snoRNA (stage 3).

## Results and discussion

Read alignments and count results are summarised in Table 1. The table shows the number and percentage of reads mapped to reference sequences [3].

**Table 1.** Summaries of reads mapping to the reference horse genome.

| Samples | #inputReads | #mappedReads | %mappedReads | #mappedPairs | %mappedPairs |
| --- | --- | --- | --- | --- | --- |
| Sample_1-eq_young | 2,879,184 | 2,711,233 | 94.17 | 1,344,271 | 93.38 |
| Sample_2-eq_young | 2,934,642 | 2,767,916 | 94.32 | 1,375,064 | 93.71 |
| Sample_3-eq_young | 2,843,524 | 2,662,410 | 93.63 | 1,312,969 | 92.35 |
| Sample_4-eq_young | 2,977,608 | 2,791,787 | 93.76 | 1,385,511 | 93.06 |
| Sample_5-eq_young | 2,464,880 | 2,268,604 | 92.04 | 1,112,517 | 90.27 |
| Sample_6-eq_old | 3,324,740 | 3,122,189 | 93.91 | 1,537,897 | 92.51 |
| Sample_7-eq_old | 3,229,210 | 3,029,631 | 93.82 | 1,493,633 | 92.51 |
| Sample_8-eq_old | 3,107,406 | 2,916,026 | 93.84 | 1,447,795 | 93.18 |
| Sample_9-eq_old | 3,246,534 | 3,023,512 | 93.13 | 1,488,651 | 91.71 |
| Sample_10-eq_old | 3,107,510 | 2,896,577 | 93.21 | 1,434,292 | 92.31 |

A total seven of novel snoRNAs in the N-S set passed all six steps of the first stage process. Six of these also passed the steps of the second stage. Figure 2 plots the cover region and coverage for novel snoRNA ECABCGRLG0000001150. It was expressed in Sample 2 (represented by the 5 black lines which are alignments of near full length), but was not evident in any other three samples. The reason is most likely that Sample_2 had far higher sequencing depth than any of other three.

**Figure 1.**
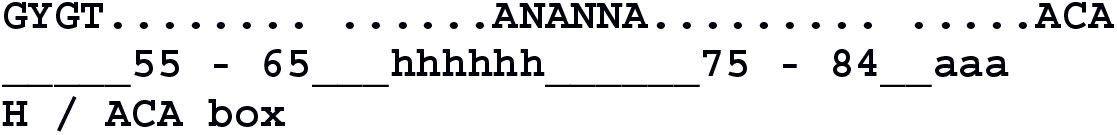
Jouvence pattern for equine. The pattern was constructed from the alignment between human and drosophila Jouvence sequences. It started with GYGT followed by 55 – 65 of any base, then a Hbox (ANANNA), followed again by 75-84 of any base, and then an ACA box.

**Figure 2.**
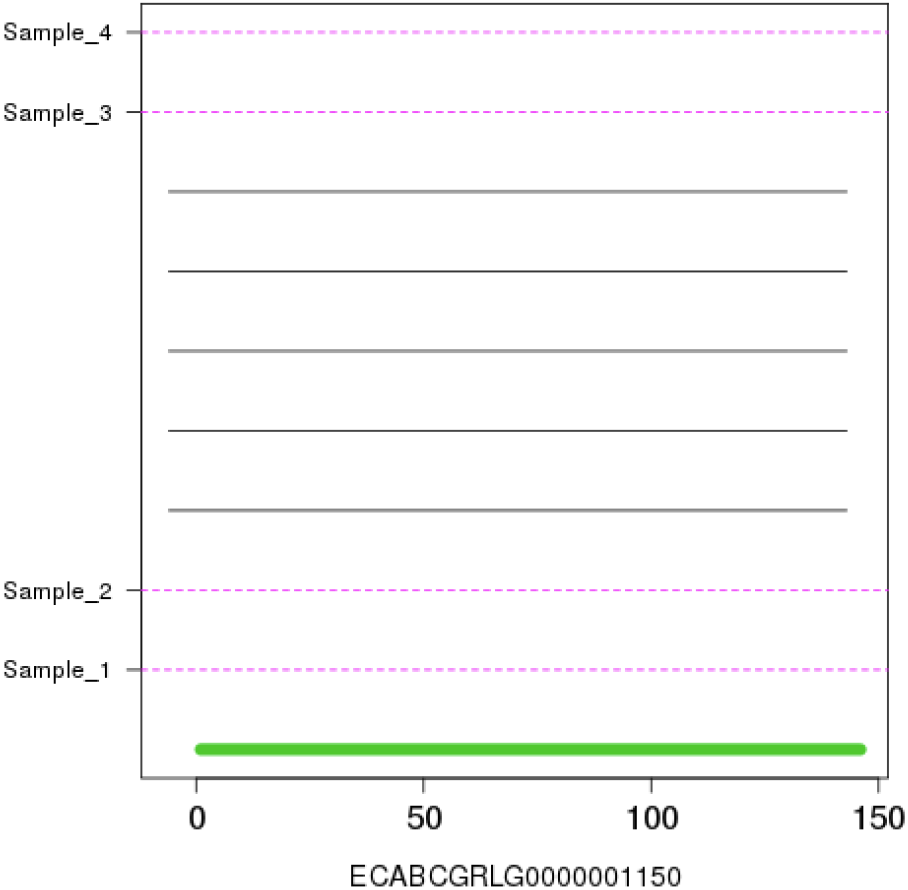
The cover region and coverage plot for novel snoRNA ECABCGRLG0000001150. The green bar represents the snoRNA feature. Besides ECABCGRLG0000000730. It was the second choice of a candidate Jouvence sequence.

After comparing to Drosophila melanogaster on secondary structure, and combining the expression information, ECABCGRLG0000000730 (internal ID) appeared a potential candidate for horse Jouvence snoRNA. Figure 3A and Figure 3B show respectively the secondary structure of Jouvence snoRNA for Drosophila melanogaster [1] and generated in this study using RNAfold [7] and for ECABCGRLG0000000730, selected from the RNAfold output [7].

**Figure 3.**
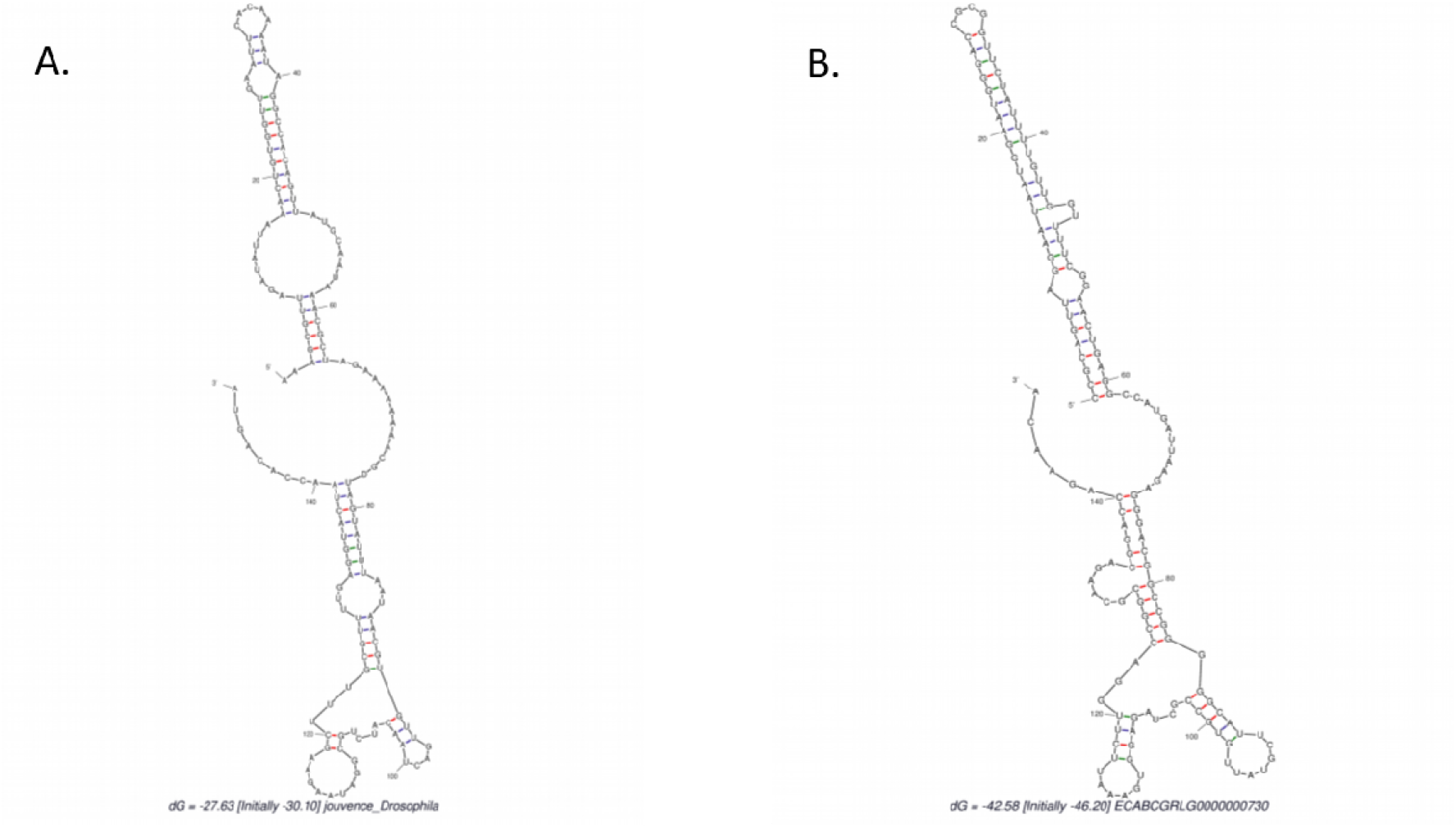
Secondary structure of Jouvence snoRNA. A. Drosophila jouvence snoRNA, B. Horse candidate Jouvence snoRNA ECABCGRLG0000000730. Images made in RNAfold [7].

For ECABCGRLG0000000730, the differential expression analysis of Jouvence snoRNA from young and old equine cartilage [3] was then undertaken (Figure 4). Although this does not indicate the effect of Jouvence snoRNA on life span it is interesting that it is significantly increased in ageing equine cartilage.

**Figure 4.**
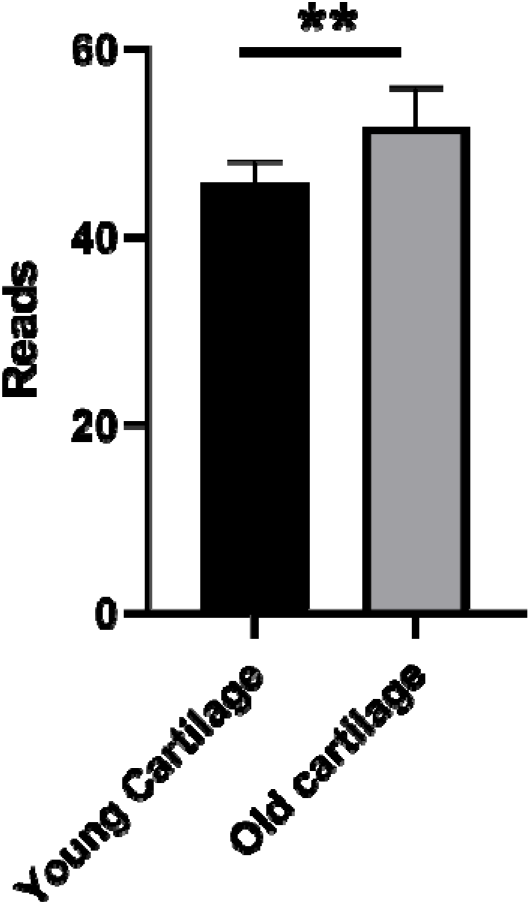
Jouvence is differentially expressed in ageing equine cartilage. Graphs shows mapped reads, individual donors and mean and standard deviation bars. Statistical Mann Whitney; P<0.01; ** plots and statistical analysis undertaken in Graphpad Prism 9.

The exploration on the P-E set sequences was carried out to answer the question of whether a better candidate horse Jouvence snoRNA exist elsewhere in the horse genome. The outcome from the P-E set did not detect any better candidate than was obtained from N-S set ECABCGRLG0000000730. In fact, after the first four steps in the first stage of processing, 21 sequences remained and only one passed the snoReport checking. None of them passed the expression check in stage 2.

## Conclusion

Putting the two investigations together, ECABCGRLG0000000730 is the primary candidate horse Jouvence snoRNA. The stage 3 process shows that the identity in pairwise global alignments is around 50%. When the secondary structures of human, mouse and horse (ECABCGRLG0000000730) were compared with the drosophila’s, ECABCGRLG0000000730 shows highest similarity with drosophila (Figure 5).

**Figure 5.**
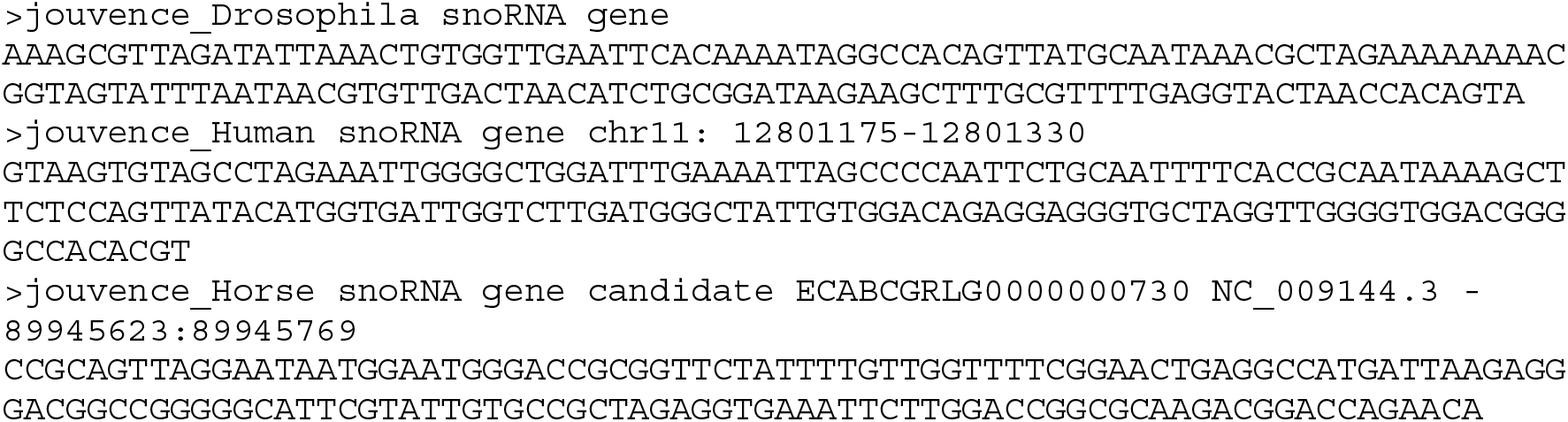
Comparison of the Jouvence snoRNA sequences for drosophila, human, mouse and proposed for horse in this study.

